# Molecular compartmentalization in a syncytium: restricted mobility of proteins within the sea urchin skeletogenic mesenchyme

**DOI:** 10.1101/2023.03.22.533866

**Authors:** Jian Ming Khor, Jennifer Guerrero-Santoro, Charles A. Ettensohn

## Abstract

Multinucleated cells, or syncytia, are found in diverse taxa. Their biological function is often associated with the compartmentalization of biochemical or cellular activities within the syncytium. How such compartments are generated and maintained is poorly understood. The sea urchin embryonic skeleton is secreted by a syncytium, and local patterns of skeletal growth are associated with distinct sub-domains of gene expression within the syncytium. For such molecular compartments to be maintained and to control local patterns of skeletal growth: 1) the mobility of TFs must be restricted to produce stable differences in the transcriptional states of nuclei within the syncytium, and 2) the mobility of biomineralization proteins must also be restricted to produce regional differences in skeletal growth patterns. To test these predictions, we expressed fluorescently-tagged forms of transcription factors and biomineralization proteins in sub-domains of the skeletogenic syncytium. We found that both classes of proteins have restricted mobility within the syncytium and identified motifs that limit their mobility. Our findings have general implications for understanding the functional and molecular compartmentalization of syncytia.

**Summary statement:** Transcription factors and effector proteins have limited mobility within the skeletogenic syncytium of the sea urchin embryo.

## Introduction

Multinucleated cells, also known as syncytia, are found across the tree of life (Ogle et al., 2005; Mela et al., 2020; Olsen, 2020; McCartney and Dudin, 2023). Syncytia can arise in two ways; through nuclear division in the absence of cytokinesis or through cell-cell fusion. In multicellular animals, syncytia first appear during embryonic development. Some embryonic syncytia, such as the cleavage-stage, syncytial embryos of many arthropods and the syncytiotrophoblast of placental mammals, are transient (Carvalho and Heisenberg, 2010; Stathopoulos and Newcomb, 2020; Renaud and Jeyarajah, 2022) while others, such as myotubes, persist into adulthood (Kim et al., 2012). Syncytia arise in adult animals during the normal process of cell differentiation, as when macrophages fuse to produce osteoclasts (Kloc et al., 2022). Syncytia can also arise under pathogenic conditions, when host cell fusion is triggered by viral and bacterial pathogens (Leroy et al., 2020; Lin et al., 2021; Bzdyl et al., 2022). In some well-studied cases, biochemical and cellular activities such as gene expression programs and patterns of nuclear division, are regionalized with syncytia (Bursztajn et al., 1989; Fogarty et al., 2011; Roberts and Gladfelter, 2013; Dundon et al., 2016; Gerber et al., 2022). Such studies reveal that distinct molecular and functional compartments can exist within a single, large, multinucleated cell. The mechanisms that underlie the generation and maintenance of such compartments, however, are poorly understood.

The development of the calcareous endoskeleton of the sea urchin embryo is a powerful model for the analysis of morphogenesis, cell differentiation, biomineralization, and the evolution of development (Oliveri et al., 2008; Ettensohn, 2009; Koga et al., 2014; McIntyre et al., 2014; McClay, 2016; Shashikant et al., 2018; Ettensohn, 2020; Gildor et al., 2021; Ben-Tabou de Leon, 2022; Ettensohn et al., 2022). The embryonic skeleton is produced by primary mesenchyme cells (PMCs), a specialized population of biomineral-forming cells derived from the micromeres of the 16 cell-stage embryo. PMCs undergo an epithelial-to-mesenchymal transition at the late blastula stage and migrate into the blastocoel cavity, where they adopt a characteristic ring-like pattern between the vegetal pole and the equator of the embryo. During this migratory phase, PMC filopodia fuse with one another, creating a slender cytoplasmic cable that connects the cells in a single, continuous, syncytial network. Amorphous calcium carbonate and associated proteins are secreted into a membrane-bound compartment within the cytoplasmic cable, and the biomineralized rods that comprise the embryonic skeleton are assembled within this compartment.

Skeletal patterning provides evidence of the regional differentiation of the PMC syncytium. Overt skeletogenesis begins at the mid-gastrula stage, when two tri-radiate skeletal rudiments are deposited within the PMC syncytium at stereotypical positions along the ventrolateral aspects of the blastocoel wall. The three arms of each skeletal rudiment subsequently elongate and branch in a stereotypical manner, with each rudiment producing an elaborate half-skeleton that is the mirror image of its partner. The different rods of the skeleton elongate at characteristic rates, and while some cease growth during embryogenesis, others grow continuously. Local variations in skeletal growth provide strong evidence of functional compartmentalization within the PMC syncytium.

Skeletal growth and patterning is mediated by short-range signals provided by the adjacent ectoderm (Okazaki, 1975; Armstrong et al., 1993; Guss and Ettensohn, 1997). Several signaling molecules secreted by the ectoderm are responsible for this control. The best characterized of these signaling molecules is VEGF3, which plays an essential role in skeletal development throughout the echinoderm phylum (Duloquin et al., 2007; Fugita et al., 2010; Knapp et al., 2012; Morino et al., 2012; Adomako-Ankomah and Ettensohn, 2013, Adomako-Ankomah and Ettensohn, 2014; Sun and Ettensohn, 2014; Ettensohn and Adomako-Ankomah, 2019; Morgulis et al., 2019; Morgulis et al., 2021). During early sea urchin embryogenesis, VEGF3 is produced by the ectoderm at the sites where the skeletal rudiments will form, and at later stages the protein is expressed by ectoderm cells at the growing tips of the skeletal rods that support the larval arms. VEGF3 interacts with a receptor tyrosine kinase, VEGFR-10-Ig, which is expressed specifically by PMCs (Duloquin et al., 2007). Two additional secreted proteins, FGF and TGFβ, also regulate skeletogenesis, although these factors are less well characterized (Röttinger et al., 2008; Sun and Ettensohn, 2017; Ankomah and Ettensohn, 2013, Adomako-Ankomah and Ettensohn, 2014). The ectodermal territories that express VEGF and FGF (and possibly TGFβ, which has not been studied in this regard) are established through the coordinated activity of several signaling pathways, including the Nodal, BMP, and Wnt pathways (Duboc et al., 2004; Flowers et al., 2004; Duloquin et al., 2007; Röttinger et al., 2008; Yaguchi et al., 2010; McIntyre et al., 2013).

PMC differentiation is controlled by a well-characterized gene regulatory network (GRN) (Oliveri et al., 2008; Shashikant et al., 2018). One cardinal function of this network is to activate the expression of a large battery of PMC-specific proteins that mediate biomineralization, including proteins that regulate calcium uptake, proton transport, bicarbonate synthesis, phase transitions of calcium carbonate, and many other proteins that affect the mineral and/or protein components of the biomineral (reviewed by Ettensohn et al., 2022). The PMC GRN network is activated prior to gastrulation in a cell-autonomous fashion within the presumptive PMCs. Based on qualitative WMISH studies, the initial, cell-autonomous phase of GRN deployment produces a homogeneous population of cells; i.e., effector genes are expressed uniformly among PMCs at the late blastula stage (Rafiq et al., 2012, 2014). During gastrulation, however, the regulation of the network shifts to a signal-dependent mode and region-specific patterns of gene expression arise within the PMC syncytium. Many mRNAs are expressed in specific sub-domains of the PMC syncytium at late developmental stages, including mRNAs that encode biomineralization proteins or transcription factors that positively regulate biomineralization genes (Harkey et al., 1992; Guss and Ettensohn, 1997; Illies et al., 2002; Livingston et al., 2006, Sun and Ettensohn, 2014). In general, regions of elevated mRNA expression correspond to regions of active skeletal growth (i.e., at the sites where the skeletal rudiments form and, at later developmental stages, at the tips of the arms). Based on these observations, it has been proposed that regional variations in skeletal growth and patterning during embryogenesis are a consequence of local, signal-dependent modulation of the skeletogenic GRN (Harkey et al., 1992; Guss and Ettensohn, 1997; Sun and Ettensohn, 2014).

Because skeletogenesis occurs within a syncytium, if localized mRNAs are to produce local differences in skeletal growth, at least two conditions must be met. First, to maintain stable, local differences in the transcriptional states of individual PMC nuclei within the syncytial network, the mobility of TFs that regulate the expression of biomineralization genes must be restricted. Rapid diffusion of these proteins would prevent the formation of stable, distinct nuclear regulatory states within the syncytium. Second, even if local differences in transcriptional regulatory states are maintained, the mobility of downstream effector proteins that directly mediate biomineralization must also be restricted to produce local control of skeletal growth. In the present study, we tested these two predictions by expressing fluorescently-tagged forms of PMC transcription factors and biomineralization proteins in sub-domains of the PMC syncytium and analyzing the mobility of the proteins in living embryos. Our findings show that both classes of proteins have restricted mobility within the PMC syncytium, providing a mechanism for the establishment and maintenance of distinct molecular and functional compartments within the syncytium.

## Materials and Methods

### Animals

Gravid, adult *Lytechinus variegatus* were obtained from Pelagic Corp (Sugarloaf Key, FL, USA). Spawning was induced by intracoelomic injection of 0.5 M KCl. *L. variegatus* embryos were cultured in artificial seawater (ASW) at 18-25°C in temperature-controlled incubators.

### DNA constructs

Plasmids used in this study were generated as previously described (Khor and Ettensohn, 2023) with a few modifications. For the transgenic activator construct, the TetOn3G recombinant gene, based on the transactivator sequence from pCAG-TetOn-3G (Faedo et al., 2017), was synthesized as a gBlock gene fragment by Integrated DNA Technologies (Coralville, IA, USA). The gBlock was cloned into EpGFPII in place of the GFP coding sequence, downstream of the *Sp-endo16* promoter. To drive PMC-specific expression, an intronic *cis*-regulatory element (CRE) of *LOC115919257* (a gene previously referred to as WHL22.691495, or *Sp-EMI/TM*, and characterized by Khor et al., 2019) was cloned upstream of the promoter to generate PMC-CRE: TetOn3G. The TRE3Gp promoter containing the Tet response element (TRE), minimal human cytomegalovirus (CMV) promoter, and CMV 5’-UTR (Kang et al., 2019) was cloned upstream of the eGFP or mCherry coding sequence to generate transgenic responder constructs. Tagged fusion proteins were generated by fusing GFP or mCherry to their C-termini with a glycine/serine-rich linker (GGGGSGGGGS).

### Doxycycline treatment

A stock solution of 10 mg/mL doxycycline hyclate (Dox) (D9891, Sigma-Aldrich: St. Louis, MO, USA) was prepared using sterile H_2_O and stored in light-protected microcentrifuge tubes at −20°C. Dox was added to the culture medium (sea water) to yield a final concentration of 5 μg/ml. In all experiments, Dox was added at the prism stage, after PMC fusion was complete and the syncytium was well-formed. Embryos were typically examined after 5-6 hours of Dox exposure, at the early two-armed pluteus stage. In some experiments, embryos were allowed to develop overnight in the presence of the drug and scored at the late two-armed pluteus stage.

### Microinjection

Linearized plasmids were injected into fertilized *L. variegatus* eggs following established protocols (Arnone et al., 2004; Cheers and Ettensohn, 2004). Each 20 μL injection solution contained 50 ng of the transactivator plasmid, 50 ng of each responder plasmid, 500 ng of HindIII-digested genomic DNA, 0.12 M KCl, 20% glycerol, and 0.1% Texas Red-Dextran (10,000 MW). Linear DNA injected into fertilized eggs forms a large concatemer that is randomly inherited by one or a few cells during cleavage (McMahon et al., 1985).

### ImmunoFISH

Combined whole-mount fluorescent in situ hybridization and immunofluorescence staining (ImmunoFISH) were carried out as previously described (Khor and Ettensohn, 2023). DNA template containing the GFP coding sequence was PCR-amplified with a reverse primer that contained a T3 promoter sequence. Digoxigenin-labeled RNA probes were synthesized using the MEGAscript T3 Transcription Kit (Invitrogen/Thermo Fisher Scientific, Waltham, MA, USA).

## Results

To examine the mobility of proteins within the PMC syncytium, we C-terminally tagged wild-type (full-length) and mutant forms of several transcription factors and biomineralization proteins with GFP or mCherry. To express proteins specifically in PMCs, we cloned their coding regions into plasmids and used a well-characterized, intronic *cis*-regulatory element (CRE) of the *S. purpuratus* gene *LOC115919257* (a gene previously referred to as WHL22.691495, or *Sp-EMI/TM*) to drive expression in transgenic embryos (Shashikant et al., 2018; Khor et al., 2019). Below, we refer to this intronic regulatory element as the “PMC CRE”. Although this CRE was originally derived from *S. purpuratus*, it also drives the PMC-specific expression of reporter genes in *L. variegatus*, the species used in this study (Khor and Ettensohn, 2023).

In sea urchins, microinjection of plasmids into 1-cell zygotes results in the mosaic incorporation and expression of transgenes (McMahon et al., 1985). In our studies, if plasmid DNA was incorporated into cells of the micromere lineage, this often produced a genetically mosaic PMC syncytium. Typically, we co-injected a plasmid encoding a complementary fluorescent protein (i.e., GFP in the case of mCherry-tagged fusion proteins and mCherry in the case of GFP-tagged fusion proteins). Many previous studies have shown that GFP and mCherry rapidly diffuse through the PMC syncytium (Arnone et al., 1997; Amore and Davidson, 2006; Wahl et al., 2009; Damle and Davidson, 2011; Shashikant et al., 2018; Khor et al., 2019; Wang et al., 2019; Khor and Ettensohn, 2022). The primary reason for co-expressing GFP or mCherry was to confirm that a complete PMC syncytium had formed in each transgenic embryos that we scored, thereby demonstrating that any restricted distribution of tagged fusion proteins we observed was a consequence of limited protein mobility within the syncytium and not a failure of cell-cell fusion. In addition, expression of GFP or mCherry provided a direct (if qualitative) comparison of the mobility of tagged fusion proteins compared to that of a highly diffusible, reference protein. The distribution of fluorescently tagged proteins was analyzed in living embryos at post-gastrula stages (i.e., after the formation of the PMC syncytium) by epifluorescence and differential interference contrast (DIC) microscopy.

In initial studies, we used a single-plasmid expression system. The coding region of an mCherry-tagged protein of interest was cloned into a modified version of the widely-used EpGFPII plasmid, with transcription of the fusion protein gene directly controlled by the PMC CRE in combination with the basal promoter of *Sp-endo16* (Fig. 1). We refer to this as a “constitutive” expression system (in contrast to an inducible system, described below), as the spatiotemporal pattern of expression of transgenes is determined solely by regulatory information intrinsic to the PMC CRE. Using this approach, we analyzed the mobility of five proteins within the PMC syncytium; three transcription factors (Sp-Alx1, Sp-Ets1, and Sp-Jun) and two biomineralization proteins (Sp-P16 and Sp-SM30B). All three transcription factors are normally expressed by PMCs during embryogenesis, and Sp-Alx1 and Sp-Ets1 provide positive regulatory inputs into many genes that have essential roles in biomineralization (Oliveri et al, 2008; Rafiq et al., 2014; Khor et al., 2019). P16 is a novel, PMC-specific transmembrane protein required for skeletal growth (Cheers and Ettensohn, 2005), and SM30B is a major protein constituent of the spicule matrix (Wilt et al., 2013). It should be noted that in these initial studies, protein-coding sequences were derived from the *S. purpuratus* genome (v.5.0) (Arshinoff et al., 2022) while microinjections were carried out using the more optically transparent eggs of *L. variegatus*. In subsequent experiments using the two-plasmid, Tet-On system (below), protein sequences were obtained from a recently improved *L. variegatus* genome assembly (Davidson et al., 2020), and expression was again assessed in *L. variegatus* embryos. In three cases (Alx1, Jun, and P16), we tested both *S. purpuratus* and *L. variegatus* forms of proteins. In these cases, we detected no species-specific differences and found that all forms were highly localized within the PMC syncytium of *L. variegatus*, as described below.

**Figure 1:**
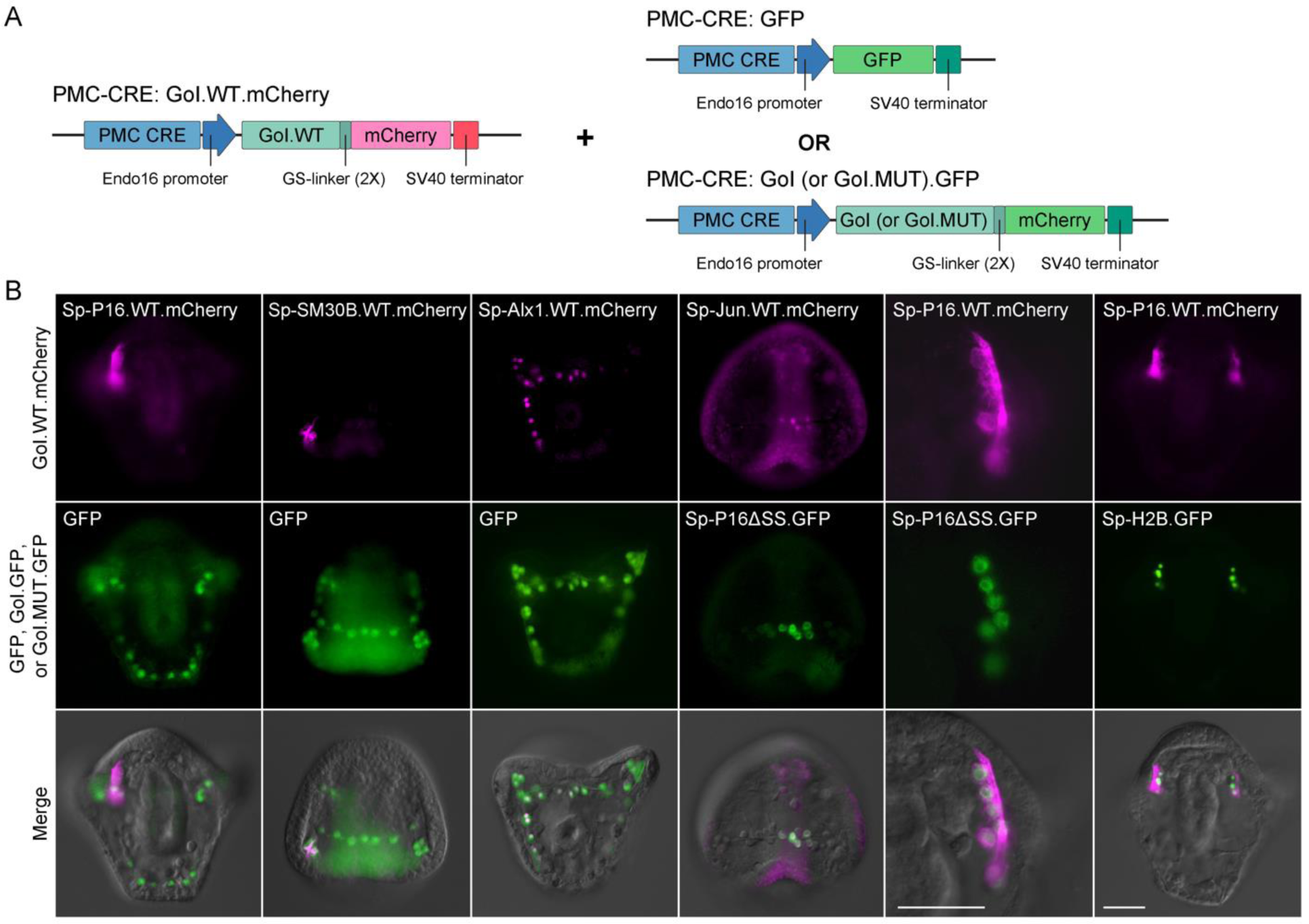
Biomineralization proteins and transcription factors exhibit restricted mobility in the PMC syncytium. (A) Schematic representation of the expression constructs used to examine protein mobility. Plasmids were based on EpGFPII and the expression of each transgene was driven specifically in PMCs by an intronic CRE of the *S. purpuratus* gene, *LOC115919257* (Shashikant et al., 2018; Khor et al., 2019) (“PMC CRE,” above). (B) Representative images of live, transgenic embryos. As GFP protein can readily diffuse throughout the PMC syncytium, the entire PMC network is labeled in transgenic embryos, despite the mosaic incorporation and expression of transgenes in sea urchins. Wild-type proteins tagged with mCherry (Sp-P16.WT.mCherry, Sp-SM30B.WT.mCherry, Sp-Alx1.WT.mCherry, and Sp-Jun.WT.mCherry) all showed localized distributions within the PMC syncytium. Deletion of the endoplasmic reticulum signal sequence of Sp-P16 (Sp-P16ΔSS.GFP) did not enhance the mobility of the protein, although the subcellular distribution of the protein was altered. Co-injection of two plasmids, each of which encoded a spatially localized protein (Sp-P16.WT.mCherry + Sp-H2B.GFP and Sp-Jun.WT.mCherry + Sp-P16ΔSS.GFP) always resulted in co-localization of the two fusion proteins, confirming that the two plasmids were expressed by the same cells. Top row: mCherry fluorescence. Middle row: GFP fluorescence. Bottom row: mCherry and GFP fluorescence overlaid onto differential interference contrast (DIC) images. GoI: Gene of Interest. Scale bar: 40 μm.

Experiments using the constitutive expression system confirmed that in 100% of the embryos examined (n >300), GFP was distributed throughout the entire PMC syncytium at post-gastrula stages (Fig. 1). GFP accumulated preferentially in PMC nuclei but was also detectable in the cytoplasm, including the PMC cytoplasmic cable. In contrast, four of the five fusion proteins exhibited a restricted distribution within the PMC syncytium (Ets1 was a special case, as described below). For each of these constructs, > 50% of transgenic embryos exhibited a restricted distribution of the fusion protein (n > 30 in all cases). Fluorescently-tagged proteins were typically expressed in a single, contiguous territory within the PMCs syncytium, even though the PMC CRE first activates gene expression prior to PMC fusion. This finding is consistent with evidence that there is little mixing of migratory PMCs prior to cell-cell fusion (Peterson and McClay, 2003). The size of the sub-domain of labeled PMCs varied from embryo to embryo, however, even within a single experimental trial. This reflected the random, mosaic incorporation and expression of transgenes in sea urchins, which would be expected to result in variable numbers of PMC progenitors (from a few to all) that expressed the constructs. Embryo-to-embryo differences in the level of expression based on position effects associated with the random insertion of the transgenes may also have contributed. In general, we observed a positive correlation between the overall intensity of the reference mCherry fluorescence and the size of the sub-domain of GFP-tagged fusion protein expression, consistent with the view that larger expression sub-domains resulted from larger clonal patches of transgenic nuclei that expressed the transgenes. We found that co-injection of two plasmids, each of which encoded a spatially localized protein (e.g., Sp-P16.WT.mCherry and a tagged histone, Sp-H2B.GFP) (Fig. 1) invariably resulted in co-localization of the two fusion proteins, confirming that the two plasmids were expressed by the same cells (Arnone et al., 1997).

The subcellular distributions of tagged fusion proteins varied. Transcription factors (Sp-Alx1.WT.mCherry and Sp-Jun.WT.mCherry) were highly concentrated in PMC nuclei (Figs. 1B, S1A). When a tagged form of Ets1 was expressed, however, we did not detect labeled nuclei within the PMC syncytium; instead, small numbers of fluorescent cells were observed in the blastocoel, unassociated with the syncytium (data not shown). This suggested that overexpression of Ets1 under the control of the PMC CRE altered the specification of presumptive PMCs. With respect to biomineralization proteins, mCherry-tagged P16 (Sp-P16.WT.mCherry) was localized on the PMC surface, including PMC filopodia and the cytoplasmic cable, while SM30B (Sp-SM30B.WT.mCherry) was concentrated in puncta, primarily in cell bodies but also along the cytoplasmic cable (Figs. 1B, S1A). Because P16 and SM30B, like many biomineralization proteins, are targeted to the secretory pathway, we asked whether the N-terminal signal sequence (SS) of P16 was required for its restricted distribution. Surprisingly, a fluorescently-tagged form of P16 that lacked this sequence (Sp-P16ΔSS.GFP) exhibited a highly restricted distribution within the PMC syncytium that appeared very similar to that of co-expressed, wild-type P16 (Fig. 1B). The subcellular distribution of P16ΔSS.GFP was distinct, however, from that of the wild-type protein. P16ΔSS.GFP was not expressed on the PMC surface but was instead retained in the cytoplasm in a reticular pattern and was concentrated in the perinuclear region (Fig 1B). This apparent subcellular targeting of P16ΔSS.GFP, perhaps attributable to the transmembrane domain of the protein, may have limited its mobility within the PMC syncytium, even in the absence of the N-terminal signal sequence.

We extended these observations using a recently developed Tet-On system for inducing transgene expression in sea urchin embryos (Khor and Ettensohn, 2023). This system utilizes two plasmids; one that carries the *reverse tetracycline-controlled transactivator (rtTA or TetOn3G)* gene under the transcriptional control of the PMC CRE and another that carries a gene encoding a fusion protein of interest under the control of Tet response elements. Transgene expression is rapidly induced when doxycycline (Dox) is added to the seawater. The inducible system offered several advantages over the constitutive system. First and most importantly, induction of protein expression after the PMC syncytium had formed provided a more direct test of the mobility of proteins that are translated within the syncytium during late embryogenesis. Second, we found that the inducible system consistently resulted in higher levels of expression of fluorescently tagged fusion proteins, which facilitated *in vivo* imaging. Lastly, we reasoned that by inducing the expression of transcription factors at late developmental stages we might avoid any re-specification of cells that resulted from overexpression of these proteins earlier in development, as appeared to occur following Ets1 overexpression.

Using this approach, we analyzed the mobility of six proteins within the PMC syncytium; three transcription factors (Lv-Alx1, Lv-Ets1, and Lv-Jun) and three biomineralization proteins (Lv-P16, Lv-SM29, and Lv-Clectin. *S. purpuratus* orthologues of four of these proteins were also examined using the constitutive expression system, as described above. The two additional proteins, Lv-SM29 and Lv-Clectin, are members of the spicule matrix protein family, which also includes SM30B (Livingston et al., 2006). Using the inducible system, wild-type or mutant forms of the proteins were tagged with GFP, and their distributions were compared to free mCherry or to other wild-type or mutant proteins tagged with mCherry. In all experiments, Dox was added at the prism stage, after PMC fusion was complete and the syncytium was well-formed. Embryos were typically examined after 5-6 hours of Dox exposure, at the early two-armed pluteus stage. For some constructs that were expressed at low levels, embryos were allowed to develop overnight in the presence of the drug and scored at the late two-armed pluteus stage.

Studies using the inducible system were consistent with those based on constitutive expression and confirmed the restricted distributions of transcription factors and biomineralization proteins within the PMC syncytium. As expected, in 100% of the embryos examined (n >400), free reporter protein (GFP or mCherry) was distributed throughout the entire PMC syncytium, over a wide range of expression levels. In contrast, each of the six fusion proteins exhibited a restricted distribution within the PMC syncytium (Fig. 2). The fraction of all embryos expressing a given transgene that exhibited a restricted domain of expression ranged from 61% to 91%, depending on the specific construct (n = 33-58). As with the constitutive expression system, fusion proteins were typically localized in a single, contiguous territory within the PMC syncytium, the size and location of which varied from embryo to embryo. The subcellular distributions of the tagged fusion proteins were consistent with those observed in studies using the constitutive system (Fig. S1). All three transcription factors (Lv-Alx1, Lv-Ets1, and Lv-Jun) were highly concentrated in PMC nuclei, while biomineralization proteins were localized on the PMC cell surface and along the spicules (Lv-P16) or in the cytoplasm and spicule compartment (Lv-SM29 and Lv-Clectin) (Figs. S1B, 2B).

**Figure 2:**
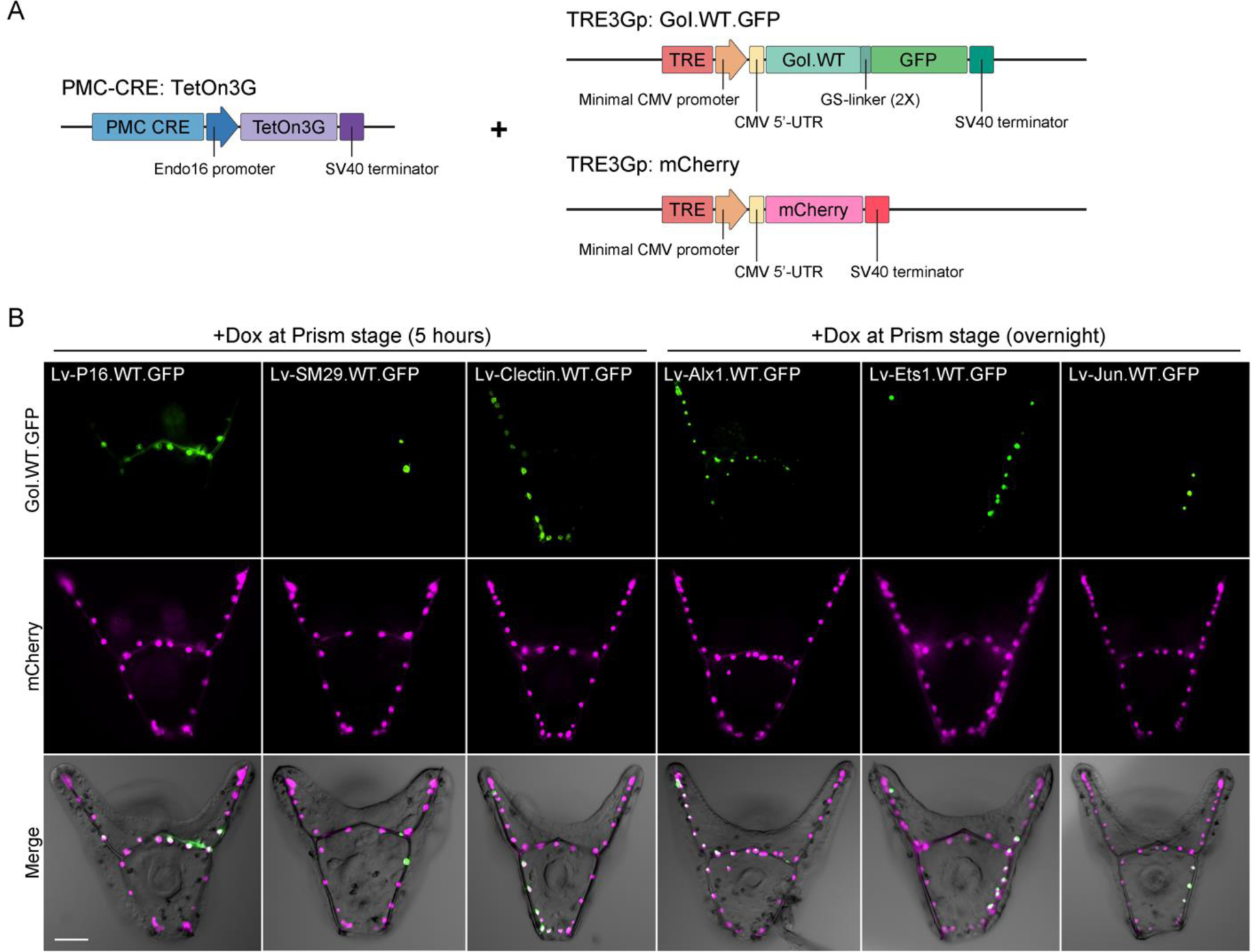
Biomineralization proteins and transcription factors exhibit restricted mobility in the PMC syncytium. (A) Schematic representation of the Tet-On transactivator and responder constructs used to induce PMC-specific expression of GFP fusion proteins and wild-type mCherry. (B) Representative images of live, transgenic embryos with Dox-induced gene expression. As mCherry protein can readily diffuse throughout the PMC syncytium, the entire PMC network is labeled in transgenic embryos, despite the mosaic incorporation and expression of transgenes in sea urchins. GFP-tagged biomineralization proteins (Lv-P16.WT.GFP, Lv-SM29.WT.GFP, and Lv-Clectin.WT.GFP) showed localized expression within the PMC syncytium. GFP-linked transcription factors (Lv-Alx1.WT.GFP, Lv-Ets1.WT.GFP, and Lv-Jun.WT.GFP) displayed strong nuclear localization in a subset of PMCs. Top row: GFP fluorescence. Middle row: mCherry fluorescence. Bottom row: GFP and mCherry fluorescence overlaid onto differential interference contrast (DIC) images. GoI: Gene of Interest. Scale bar: 50 μm.

Theoretically, the localization of tagged proteins in specific subdomains of the PMC syncytium could arise through long-distance protein translocation coupled with selective targeting to (or trapping at) specific sites. This mechanism seemed extremely unlikely given that every protein we tested exhibited a restricted distribution, and in each case we observed diverse patterns of localization within the syncytium. Nevertheless, to directly test whether proteins translocate over long distances or instead remain near the site of translation, we carried out combined immunofluorescence/RNA in situ hybridization on transgenic embryos that expressed Lv-Alx1.GFP. The distribution of *Lv-alx1.GFP* mRNA was assessed by in situ hybridization using a labeled probe complementary to the GFP coding sequence, and the distribution of Lv-Alx1.GFP protein was examined in the same specimen using an affinity-purified antibody that recognizes GFP. These experiments revealed that reporter mRNA and protein were co-localized in the same sub-domain of the PMC syncytium, with the distribution of the Lv-Alx1.GFP protein slightly broader than that of the mRNA (Fig. 3). We conclude that the restricted distribution of Lv-Alx1.GFP, and presumably that of other proteins, is not due to long-distance targeting or trapping, but instead reflects the tendency of proteins to remain near the site of synthesis.

**Figure 3:**
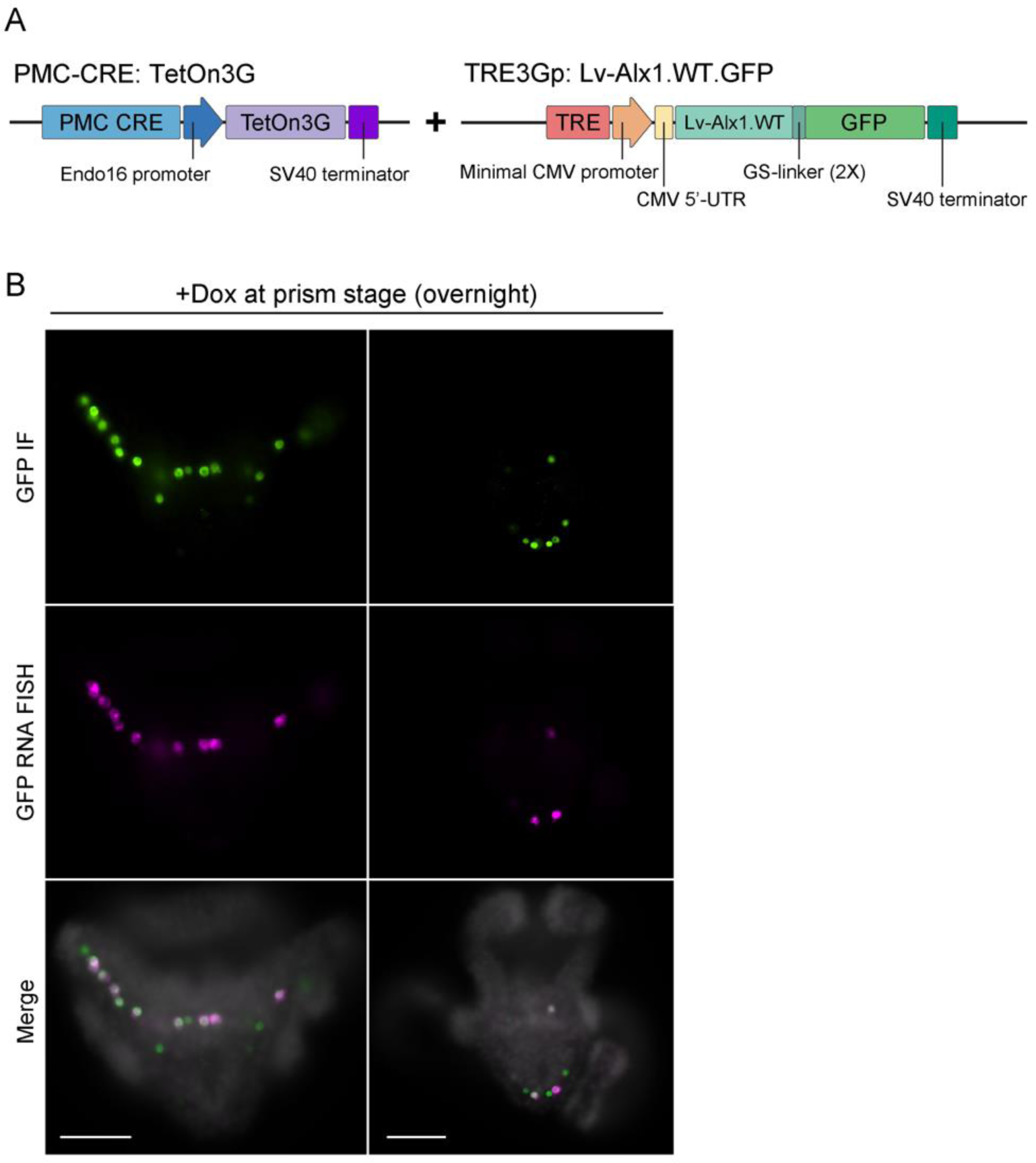
The localization of proteins within the PMC syncytium in transgenic embryos is due to limited mobility and not long-distance trafficking. (A) Schematic representation of the Tet-On transactivator and responder constructs used to induce PMC-specific expression of Lv-Alx1.WT.GFP. (B) GFP immunoFISH labeling of two fixed, transgenic embryos expressing Lv-Alx1.WT.GFP. In each embryo, Lv-Alx1.WT.GFP protein was localized to the same sub-domain of the PMC syncytium as *Lv-alx1.WT.GFP* mRNA. In each case, the distribution of the GFP-tagged fusion protein was slightly broader than that of the cognate mRNA. Top row: GFP-immunostained cells. Middle row: Cy3-labeled *gfp* RNA transcripts. Bottom row: Fluorescence merged with Hoechst 33342 counterstain (shown in grayscale). Scale bar: 50 μm.

We next analyzed the roles of specific protein domains in restricting mobility within the PMC syncytium. Each transcription factor we examined contains a DNA-binding domain flanked by nuclear localization sequences (Boulukos et al., 1989; Tagawa et al., 1995; Okamura et al., 2009). Deletion of the complete DNA binding domains (including the flanking nuclear localization sequences) of Lv-Alx1, Lv-Ets1, and Lv-Jun dramatically increased the mobility of all three proteins within the syncytium relative to the corresponding, wild-type forms (Fig. 4). The mutant forms of these proteins were also less tightly restricted to the nucleus than the wild-type forms (Fig. 4B, column on right). Next, we tested whether the DNA binding domains of Alx1, Ets1, and were sufficient to restrict protein mobility when fused to fluorescent reporters. Through co-expression studies, we directly compared the mobility of reporter proteins fused to the complete DNA binding domains of each of the three transcription factors (Lv-Alx1.DBD.mCherry, Lv-Ets1.DBD.mCherry, and Lv-Jun.DBD.mCherry) to the mobility of reporter proteins fused to DNA binding domains that lacked flanking nuclear localization motifs (Lv-Alx1.DBDΔNLS.GFP, Lv-Ets1.DBDΔNLS.GFP, and Lv-Jun.DBDΔNLS.GFP) (Fig. 5). The results of these studies were variable depending on the specific construct. In general, fusion to DNA binding domains (with or without flanking nuclear localization sequences) did not restrict the mobility of fluorescent reporters within the PMC syncytium to the extent seen when reporters were fused to wild-type, full-length proteins (compare Figs. 2 and 5). Some partial effects were seen, however, particularly in the case of the complete DNA binding domain of Lv-Ets1, which markedly reduced the mobility of mCherry.

**Figure 4:**
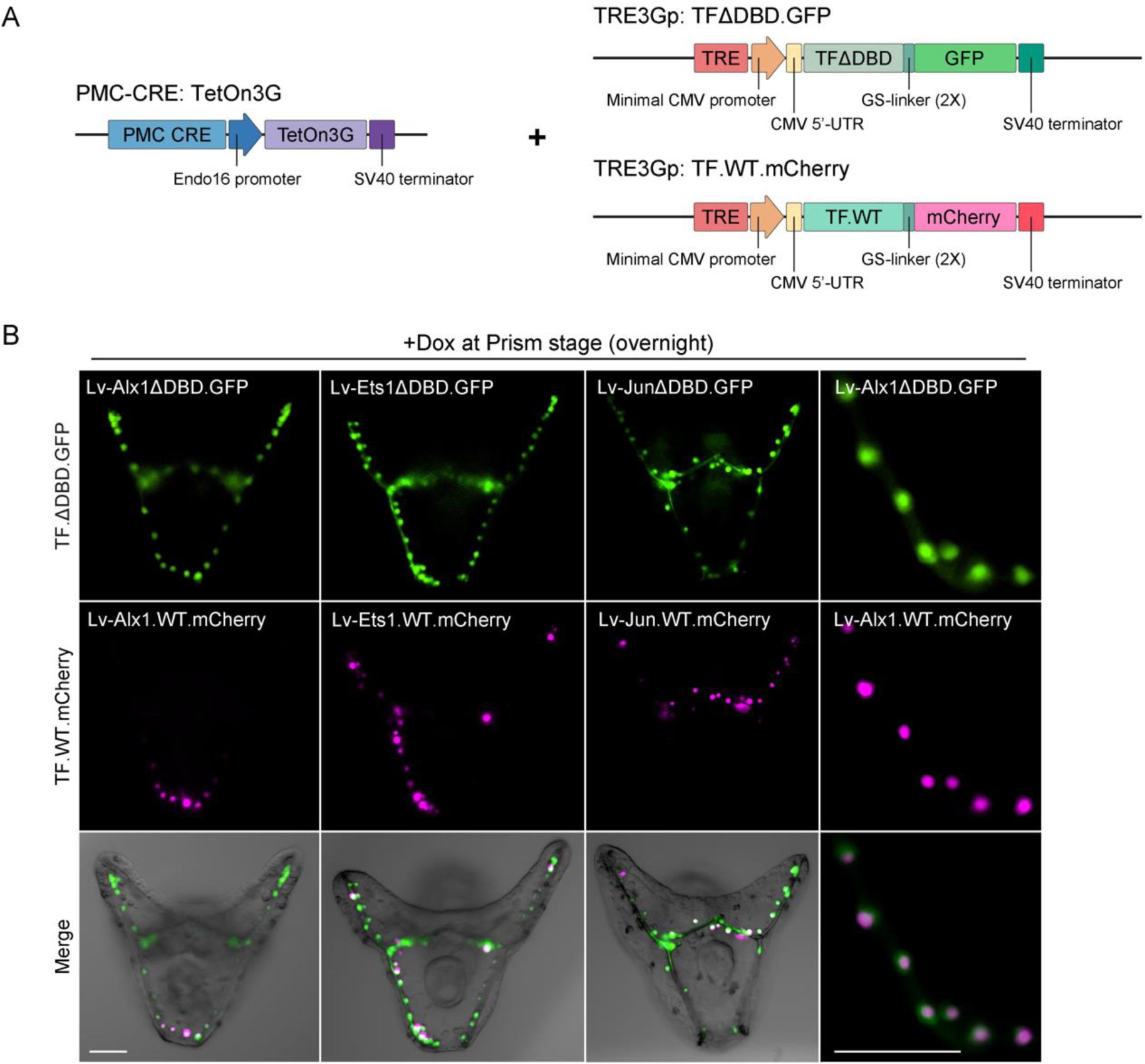
Sea urchin transcription factors lacking DNA-binding domains (DBDs) readily diffuse throughout the PMC syncytium. (A) Schematic representation of the Tet-On transactivator and responder constructs used to induce PMC-specific expression of GFP and mCherry fusion proteins. (B) Representative images of live, transgenic embryos after Dox-induced gene expression. GFP-linked transcription factors lacking DBDs (Lv-Alx1ΔDBD.GFP, Lv-Ets1ΔDBD.GFP, and Lv-JunΔDBD.GFP) were distributed much more widely within the syncytium than their wild-type counterparts (Lv-Alx1.WT.mCherry, Lv-Ets1.WT.mCherry, and Lv-Jun.WT.mCherry). Deletion of DBDs also caused transcription factors to be less tightly restricted to the cell nucleus, as shown here for Lv-Alx1. Top row: GFP fluorescence. Middle row: mCherry fluorescence. Bottom row: GFP and mCherry fluorescence overlaid onto differential interference contrast (DIC) images. Scale bars: 50 μm.

**Figure 5:**
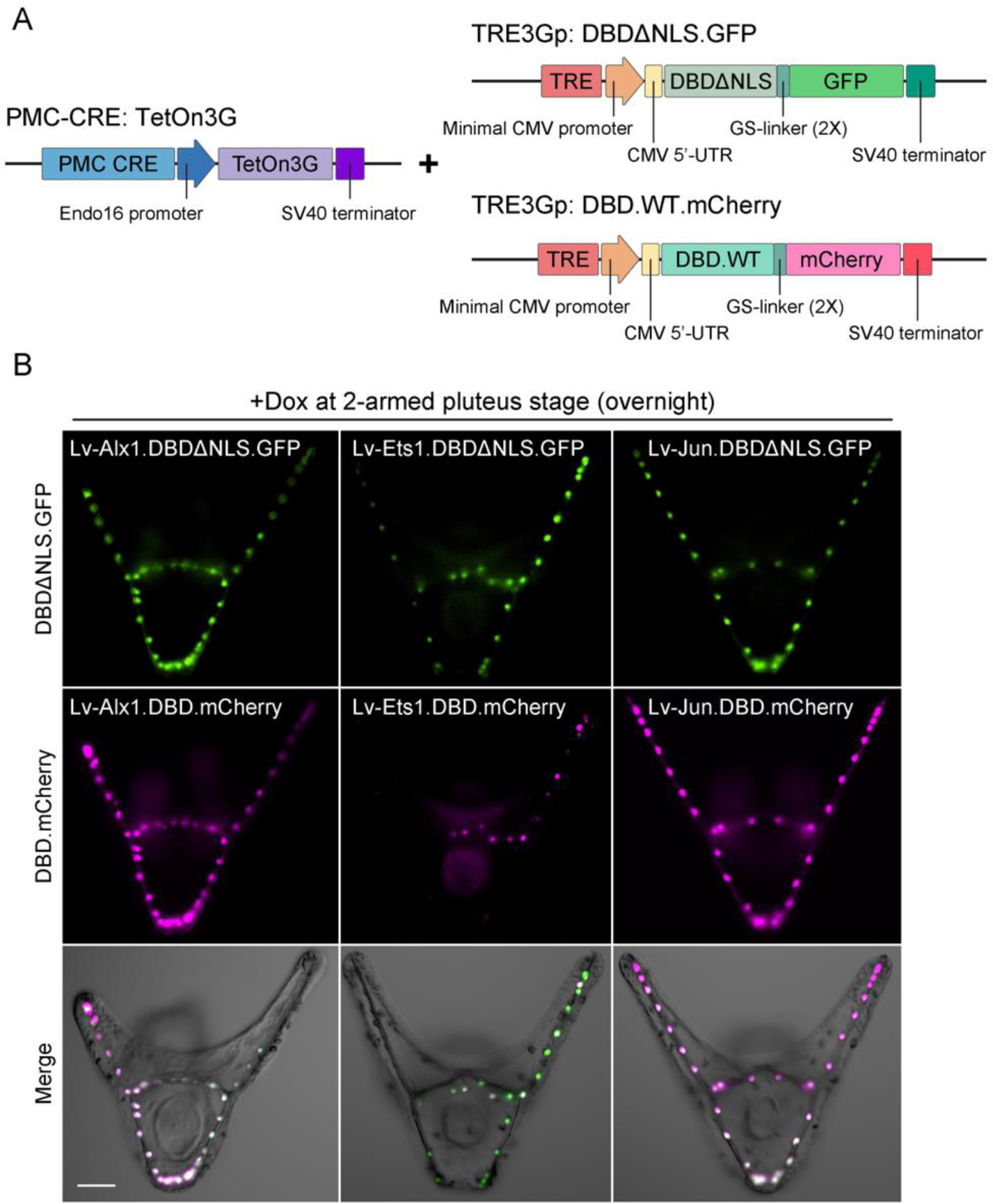
The DNA-binding domain (DBD) of Lv-Ets1, but not those of other transcription factors expressed by PMCs, is sufficient to reduce the mobility of mCherry. (A) Schematic representation of the Tet-On transactivator and responder constructs used to induce PMC-specific expression of GFP and mCherry fusion proteins. Fusion proteins containing the complete DBDs (DNA-binding motifs + nuclear localization sequences) of Lv-Alx1 (Lv-Alx1.DBD.mCherry) and Lv-Jun (Lv-Jun.DBD.mCherry) were distributed throughout the PMC syncytium, as were fusion proteins containing DBDs that lacked nuclear localization sequences (Lv-Alx1.DBDΔNLS.GFP and Lv-Jun.DBDΔNLS.GFP). In contrast, fusion of the complete DBD of Lv-Ets1 to mCherry (Lv-Ets1.DBD-mCherry) markedly reduced the mobility of the protein. This effect was dependent, at least in part, on nuclear localization sequences within the DBD, as shown by the increased mobility of Lv-Ets1.DBDΔNLS.GFP. Top row: GFP fluorescence. Middle row: mCherry fluorescence. Bottom row: GFP and mCherry fluorescence overlaid onto differential interference contrast (DIC) images. Scale bar: 50 μm.

Many proteins that mediate biomineralization are targeted to the spicule compartment via the secretory pathway (Livingston et al., 2006; Mann et al., 2010). We hypothesized that secretory activity within the PMC syncytium might be localized, thereby restricting the distribution of such proteins. To test this hypothesis, we examined the distribution of mutant forms of Lv-P16, Lv-SM29, and Lv-Clectin that lacked N-terminal, endoplasmic reticulum (ER) signal sequences (Fig. 6). Deletion of the signal sequence of Lv-Clectin (Lv-ClectinΔSS.GFP) dramatically expanded the distribution of the protein compared to the wild-type form. Deletion of the N-terminal signal sequence of Lv-SM29 also expanded the distribution of the protein within the syncytium but more modestly; the mutant protein remained enriched near the site of synthesis based on the overlapping distribution of wild-type Lv-SM29. Wild-type Lv-SM29 was localized in prominent puncta that were often observed in filopodia distant from PMC cell bodies (Fig. S1B), while the mutant form was localized in PMC cell bodies and was not concentrated in puncta. In contrast, and consistent with observations made using the constitutive expression system, deletion of the signal sequence of Lv-P16 (Lv-P16ΔSS.GFP) had no detectable effect on the distribution of the protein within the PMC syncytium, although it altered the subcellular localization of the protein, as described above (Fig. 1B). P16 also contains a predicted transmembrane domain, which is located near the C-terminus of the protein. Deletion of the transmembrane domain alone (Lv-P16ΔTM.GFP) also had little or no effect on the distribution of the protein (Fig. 6B). Deletion of both the N-terminal signal sequence and the transmembrane domain, however, greatly enhanced the mobility of the protein within the PMC syncytium. This mutant form of Lv-P16 was localized primarily in the cytoplasm.

**Figure 6:**
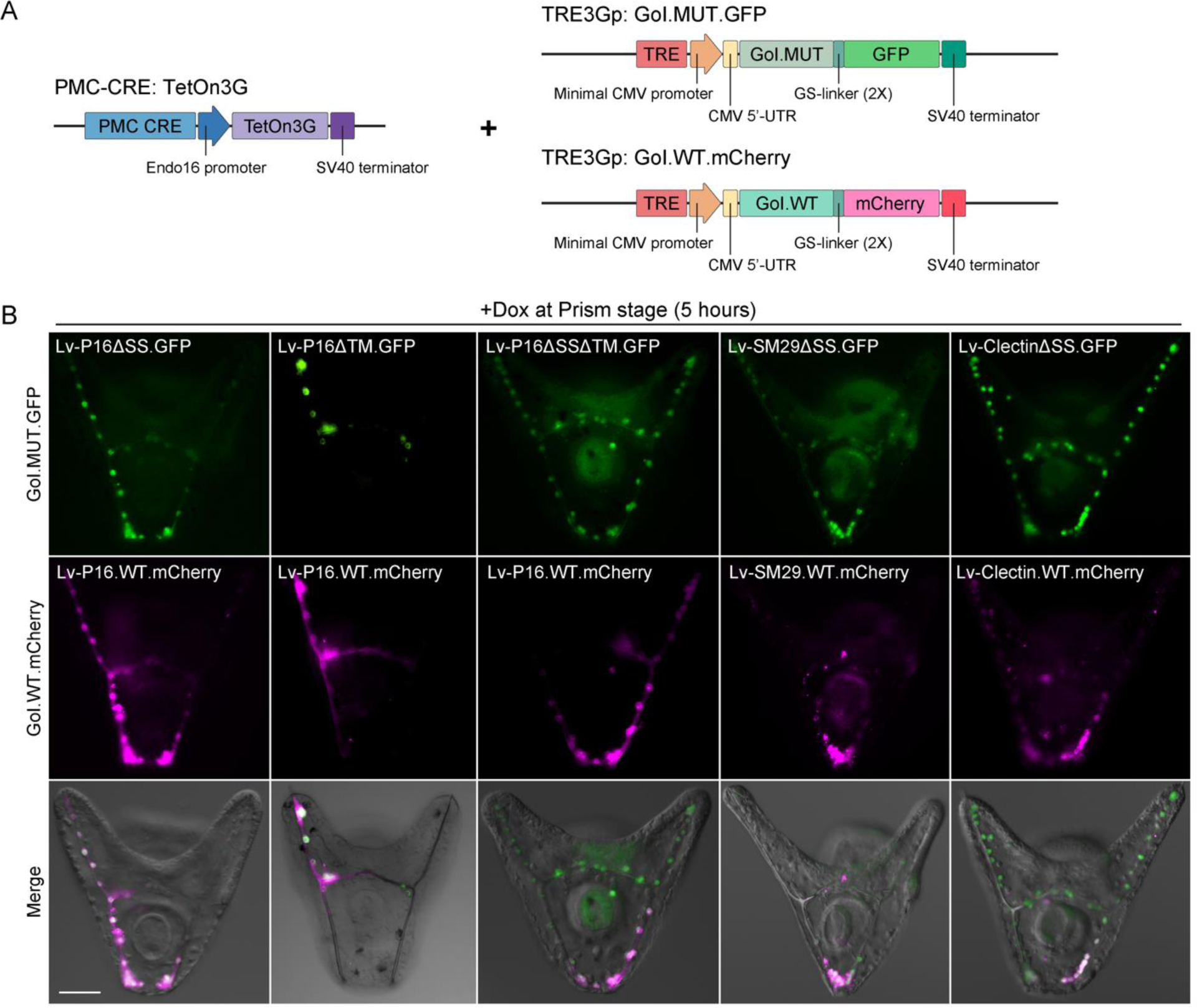
Deletion of targeting sequences from sea urchin biomineralization proteins increases their mobility within the PMC syncytium. (A) Schematic representation of the Tet-On transactivator and responder constructs used to induce PMC-specific expression of GFP and mCherry fusion proteins. (B) Representative images of live, transgenic embryos with Dox-induced gene expression. Single deletions of the N-terminal endoplasmic reticulum signal sequence or the C-terminal transmembrane domain of Lv-P16 (Lv-P16ΔSS.GFP had Lv-P16ΔTM.GFP) had little effect on the distribution of the protein. Deletion of both motifs, however (Lv-P16ΔSSΔTM.GFP), markedly increased mobility. Deletion of the signal sequences of Lv-Clectin (Lv-ClectinΔSS.GFP) and Lv-SM29 (Lv-SM29ΔSS.GFP), proteins that lack other targeting motifs, increased mobility relative to the corresponding, wild-type forms of these proteins, although this effect was more pronounced in the case of Lv-Clectin. Top row: GFP fluorescence. Middle row: mCherry fluorescence. Bottom row: GFP and mCherry fluorescence overlaid onto differential interference contrast (DIC) images. GoI: Gene of Interest. Scale bar: 50 μm.

To further explore the role of N-terminal signal sequences in regulating the mobility of proteins within the PMC syncytium, we examined the distribution of chimeric forms of GFP that contained the N-terminal signal sequences of Lv-P16, Lv-SM29, or Lv-Clectin (Fig. 7). In all cases, N-terminal signal sequences were inserted immediately downstream of the start codon and were separated from the remainder of the GFP polypeptide by two tandem copies of a flexible, glycine/serine-rich linker. Each of these mutant forms of GFP was detected only at very low levels in the PMC syncytium, distinctly lower than when GFP was expressed alone, consistent with observations made using the constitutive expression system (data not shown). This may have been because the folding and stability of GFP was altered by addition of the N-terminal signal sequence or because the fusion protein was secreted into the extracellular environment. Others have reported that GFP folding can be perturbed when fusion proteins are targeted to the lumen of the ER (Valbuena et al., 2020). Nevertheless, in embryos with detectable levels of these fusion proteins, the proteins were frequently restricted to a sub-domain of the PMC syncytium (>30% of embryos in each case), which was never observed with GFP. The subcellular distribution of the fusion proteins was distinct from that of wild-type Lv-P16, Lv-SM29 and Lv-Clectin, as the former showed relatively homogeneous cytoplasmic localization and were not found in puncta. As P16 contains a transmembrane domain in addition to an N-terminal signal sequence, and because our previous studies suggested that the transmembrane domain might influence the mobility of P16 within the syncytium, we also tested the effect of linking the Lv-P16 transmembrane domain to GFP. As in studies with N-terminal signal sequences, the Lv-P16 transmembrane domain was inserted immediately downstream of the start codon and separated from the remainder of the GFP sequence by two tandem copies of a glycine/serine-rich spacer. The chimeric Lv-P16TM.GFP protein exhibited a restricted distribution in 87% of transgenic embryos that expressed the protein (n = 23) (Fig. 7), indicating that the P16 transmembrane sequence was sufficient to limit protein mobility within the PMC syncytium. A modified form of this protein that also contained the N-terminal signal sequence of Lv-P16 exhibited a similar, highly restricted distribution (Fig. 7).

**Figure 7:**
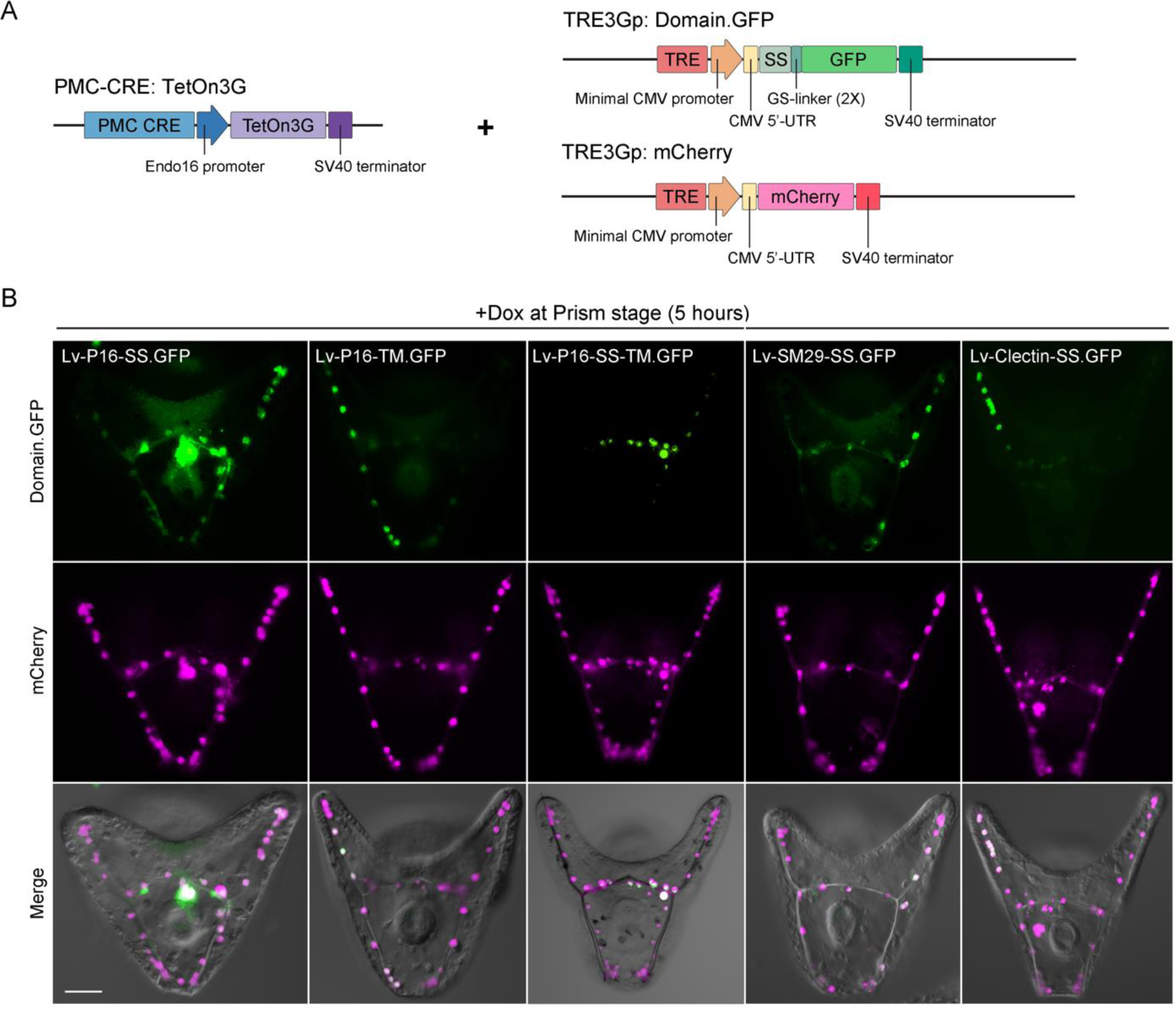
Fusion of targeting sequences of sea urchin biomineralization proteins to GFP reduces mobility. (A) Schematic representation of the Tet-On transactivator and responder constructs used to induce PMC-specific expression of GFP fusion proteins and wild-type mCherry. (B) Representative images of live, transgenic embryos with Dox-induced gene expression. N-terminal fusions of the signal sequences (SS) of Lv-SM29, Lv-Clectin, or Lv-P16 to GFP (Lv-P16-SS.GFP, Lv-SM29-SS.GFP, and Lv-Clectin-SS.GFP), or fusion of the transmembrane domain of P16, alone (Lv-P16-TM.GFP) or in combination with the Lv-P16 signal sequence (Lv-P16-SS-TM.GFP) reduced the mobility of the fusion proteins within the PMC syncytium compared to mCherry. Top row: GFP fluorescence. Middle row: mCherry fluorescence. Bottom row: GFP and mCherry fluorescence overlaid onto differential interference contrast (DIC) images. Scale bar: 50 μm.

## Discussion

We have used the skeletogenic mesenchyme of the sea urchin embryo as a model for discovering mechanisms that generate specialized molecular and functional domains within syncytia. Our studies show that the mobility of transcription factors and biomineralization proteins is restricted within the PMC syncytium. The mobility of proteins within the PMC syncytium is likely influenced by several factors. Protein size alone is evidently not a major determinant of mobility, as we have documented several relatively large proteins (MW = 66-76 kDa) that are highly mobile, including Lv-Alx1ΔDBD.GFP, Lv-Ets1ΔDBD.GFP (both from this study), Sp-caMEK.GFP, and Sp-DUSP6.GFP (Khor and Ettensohn, 2023), as well as much smaller proteins with very limited mobility (e.g., Lv-P16TM.GFP, MW = 31.5 kDa). Many of the proteins that regulate biomineralization, including those of the MSP130, P16, and spicule matrix protein families, are secreted or membrane-associated proteins (Livingston et al, 2006; Mann et al., 2010). The presence of N-terminal, ER signal sequences on these proteins could recruit these proteins co-translationally to nearby ER membranes and thereby locally target secretion. Notably, ultrastructural analysis of the PMC syncytium has shown that Golgi stacks are restricted to the perinuclear regions of PMC cell bodies and are completely absent from the cytoplasmic cable (Gibbins et al., 1969), suggesting that secretory activity is spatially localized within the syncytium at the level of individual cell bodies. In the present study, we found that N-terminal fusions of the ER signal sequences of biomineralization proteins markedly reduced the mobility of GFP within the syncytium. In addition, deletion of the N-terminal ER signal sequences of biomineralization proteins expanded their distributions within the syncytium, although to varying degrees, and the mobility of one protein (P16) was also regulated by its transmembrane domain. With respect to transcription factors, deletion of DNA binding domains and their associated nuclear localization sequences demonstrated the importance of these sequences in restricting mobility within the syncytium. Fusions of these sequences to fluorescent reporters, however, suggested that other domains of transcription factors also contribute to their restricted mobility. These might include, for example, motifs that mediate protein-protein interactions.

It is evident that the distributions of exogenous, tagged proteins described in this study do not directly reflect the distributions of the corresponding, endogenous proteins. First, the levels of expression of the exogenous and endogenous proteins are almost certainly different. Second, endogenous mRNAs are often more tightly localized within the PMC syncytium than are the clonal territories of mRNA expression we observed in transgenic embryos, implying that endogenous proteins can have much more restricted distributions than the patterns of exogenous reporters we observed. For example, at late embryonic stages, *p16* mRNA is restricted to just 2-3 cells at the tip of each arm (Cheers and Ettensohn, 2005). Lastly, many (although not all) genes in the skeletogenic GRN are expressed by all PMCs early in development and only later show restricted expression in sub-domains of the syncytium. The extent to which initial, broad patterns of protein expression are replaced by region-specific patterns will depend on the perdurance of the proteins synthesized early in development. Taken together, these considerations suggest that the distributions of endogenous proteins within the PMC syncytium depend on a complex combination of factors, including RNA expression patterns, RNA and protein expression levels, protein stability, and the mobility of proteins within the syncytium. Our findings support the view, however, that restricted protein mobility is one important factor in establishing the heterogeneous distributions of SM30 (Urry et al., 2000), Tbx2/3 (Gross et al, 2003), and phospho-SMAD1/5/8 (Luo and Su, 2012), which have been documented using antibodies against the endogenous proteins.

This study does not address the mechanisms that underlie the localized distributions of mRNAs within the PMC syncytium. The highly localized expression in transgenic embryos of a wide variety of experimentally expressed mRNAs, which lack any native, subcellular targeting motifs, clearly shows that exogenous RNAs have very limited mobility within the PMC syncytium. We strongly favor the view that the same is true of endogenous mRNAs; i.e., that mRNAs remain near the site of synthesis. Formally, however, the possibility cannot be excluded that endogenous mRNAs translocate over long distances and are directionally targeted to (or trapped at) specific sites within the PMC syncytium. Complex mechanisms would be required, however to account for the long-distance movements of a large number of endogenous mRNAs and their selective targeting to several diverse locations within the syncytium.

Our findings are consistent with a model of skeletal patterning which proposes that local, ectoderm-derived cues control the expression or activity of regulatory (i.e., transcription factor-encoding) genes in the skeletogenic GRN, creating sub-domains of gene expression within the syncytium that are stable due to the limited mobility of transcription factors (Fig. 8). According to this model, local differences in the regulatory states of PMC nuclei subsequently lead to differences in the expression of downstream effector genes, including those that are regulators of biomineral growth. Given the complex suite of signaling molecules that regulates skeletal growth, there may be diverse signaling environments within the syncytium that generate multiple, distinct regulatory states. This could contribute to the complex, overlapping expression patterns of many biomineralization genes controlled by the PMC GRN (Sun and Ettensohn, 2014). The limited mobility of biomineralization proteins within the syncytium would be expected to generate sub-domains characterized by distinct constellations of biomineralization proteins, which could then lead to local patterns of skeletal growth. Beyond simply determining whether biomineral is deposited or not in a given sub-domain of the syncytium (as illustrated in the simplified model shown in Fig. 8), we hypothesize that local expression patterns of biomineralization proteins regulate more subtle aspects of skeletal patterning, such as skeletal rod morphology (simple versus fenestrated, smooth versus barbed), growth rate, crystallographic axis of growth, and sites of branching, all of which are features of growth that could be regulated by proteins associated with the growing biomineral (Weiner, 2008; Veis, 2011; Marin et al., 2018).

**Figure 8:**
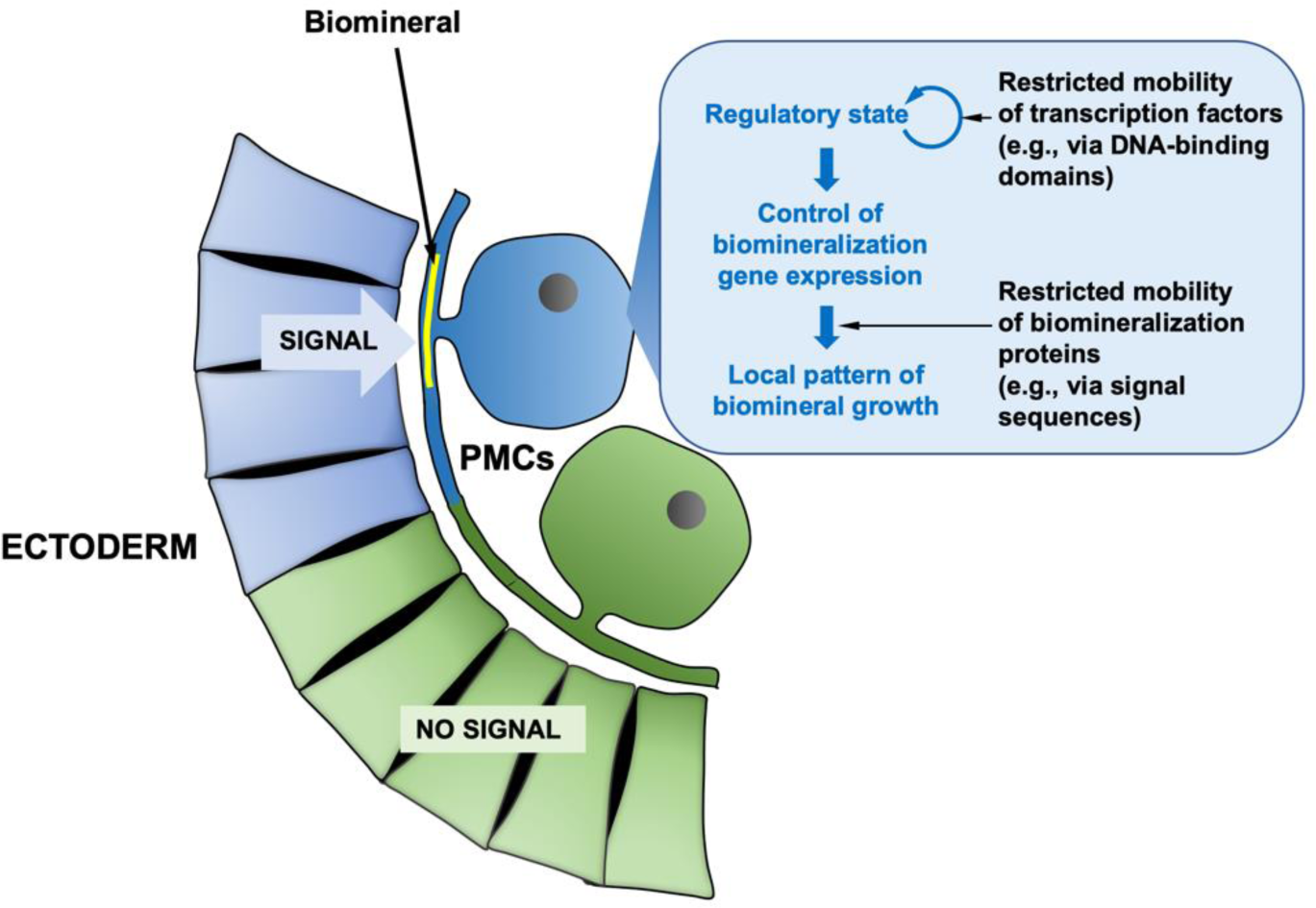
A model for the local control of skeletal growth in the PMC syncytium. Signaling ligands, including VEGF3, are secreted by specific ectodermal territories and lead to the localized expression of specific regulatory (i.e., transcription factor-encoding) genes in the nuclei of nearby PMC cell bodies. Distinct nuclear regulatory states are maintained in sub-domains of the syncytium due to the limited mobility of transcription factors. Local regulatory states, in turn, lead to territory-specific expression of downstream effector genes, including genes that encode key regulators of biomineral growth. Biomineralization proteins also have limited mobility, creating sub-domains of the syncytium with distinct assemblages of biomineralization proteins. Region-specific patterns of biomineralization proteins then lead to local patterns of skeletal growth. Note that the diagram illustrates only two distinct nuclear regulatory states (blue and green), while there may be many diverse regulatory stages within the PMC syncytium. The diagram is also simplified as it suggests that signaling determines only whether or not biomineral is deposited, while we hypothesize that region-specific expression of biomineralization proteins also regulate other, more subtle, aspects of skeletal patterning, such as skeletal rod morphology, growth rate, and branching (see Discussion). Also note that the model requires that one or more essential components of signal transduction pathways downstream of PMC receptors have limited mobility within the PMC syncytium, although this is not shown.

Although this model emphasizes regulation at the transcriptional level, post-translational mechanisms may also contribute to local patterns of skeletal growth. Here again, the restricted mobility of proteins within the syncytium is likely to play a key role. If ectoderm-derived cues result in the post-translational modification of PMC transcription factors, the inherent, limited mobility of transcription factors within the syncytium (which we observed with all transcription factors tested) would generate sub-domains of activated (or repressed) forms. Considerable evidence supports the view that such a mechanism operates during normal skeletogenesis and accounts for the regulation of skeletal growth by VEGF3. A key transcription factor in the PMC GRN, Ets1, is positively regulated by ERK signaling (Röttinger et al., 2004), a pathway activated by VEGF receptors in many cell types (Cross et al., 2003; Holmes et al., 2007; Claesson-Welsh and Welsh, 2013). Although the pivotal role of Ets1 in the initial, cell-autonomous activation of the skeletogenic GRN is well-established, until recently it was unknown whether Ets1 continues to provide regulatory inputs at post-gastrula stages, when the PMC GRN and skeletal growth are regulated by VEGF3. Recent studies using the conditional, Tet-On system to perturb Ets1 function and manipulate the expression of positive and negative regulators of the ERK pathway specifically in PMCs have shown that Ets1 and ERK signaling continue to regulate skeletogenesis after the formation of the PMC syncytium (Khor and Ettensohn, 2023). The restricted mobility of Ets1, the regulation of its activity by ERK signaling, and the essential late inputs from Ets1 and ERK into skeletogenic genes, make Ets1 an attractive candidate for linking ectodermal signaling to skeletal patterning.

Our findings are relevant to the functional and molecular compartmentalization of other syncytia. Heterogeneous patterns of gene expression have been documented in several other syncytia, including vertebrate muscle (Bursztajn et al., 1989) and syncytiotrophoblast (Fogarty et al., 2011), slime mold pseudoplasmodia (Gerber et al., 2022), and multinucleate fungi (Dundon et al., 2016), although the molecular mechanisms that establish and maintain these domains are not well understood. In the best-studied case, the early, syncytial development of Drosophila embryos, anterior-posterior compartments of gene expression are generated through the limited diffusibility of several transcription factors, including Bicoid (Huang and Saunders, 2020). The restricted mobility of transcription factors is likely to be a pre-requisite for the establishment of gene expression compartments in syncytia. In some instances, such as the skeletogenic syncytium of the sea urchin embryo, the limited mobility of effector proteins encoded by downstream target genes then generates functional compartments within the syncytium.

## Acknowledgements

The authors are grateful to Dr. Brooke McCartney for many valuable suggestions that improved the manuscript. This work was supported by National Science Foundation grant IOS2004952 (C.A.E).

---

**Figure S1:**
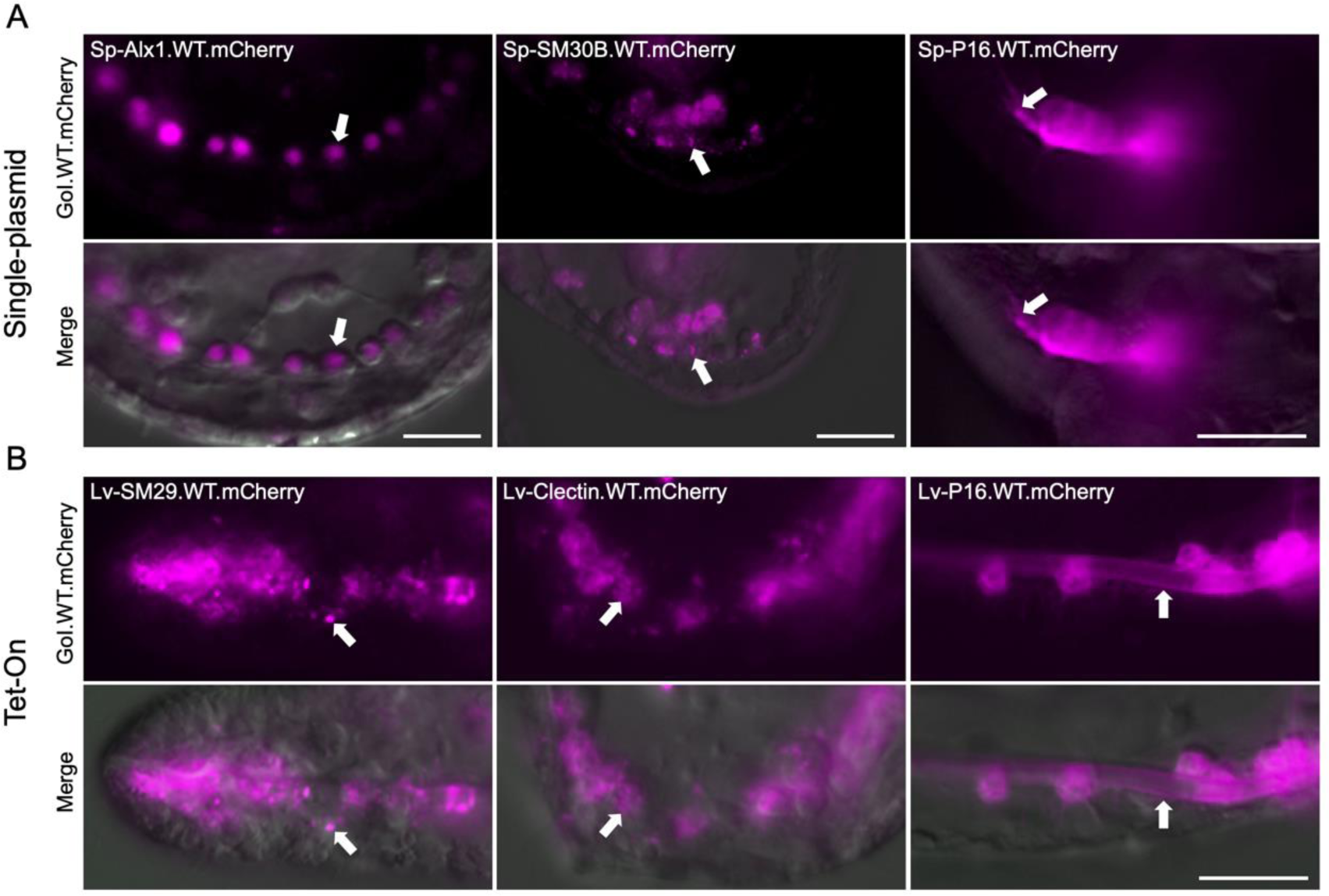
High magnification views illustrating the subcellular distributions of wild-type, mCherry-tagged proteins in PMCs. All images show live embryos. (A) Transgene expression driven using the constitutive (single-plasmid) system. Transcription factors, including Sp-Alx1, are concentrated in PMC nuclei (arrows). The spicule matrix protein Sp-SM30B is found in puncta in cell bodies (arrows) and cytoplasmic extensions. Sp-P16 is expressed on the PMC surface, including the cytoplasmic cable and along PMC filopodia (arrows). Top row: mCherry fluorescence. Bottom row: mCherry fluorescence overlaid onto differential interference contrast (DIC) images. Scale bars = 15 μm. (B) Transgene expression driven by the inducible (Tet-On) system. The spicule matrix proteins Lv-SM29 and Lv-Clectin, like Sp-SM30B, are concentrated in puncta in cell bodies and cytoplasmic projections (arrows). Lv-P16 is concentrated on the PMC surface, including the surface of the cytoplasmic cable that contains the growing biomineral (arrow). Top row: mCherry fluorescence. Bottom row: mCherry fluorescence overlaid onto differential interference contrast (DIC) images Scale bar: 15 μm.

## Notes

### Competing Interest Statement

The authors have declared no competing interest.

